# The effect of sex on sleep EEG in typical and altered neurodevelopment

**DOI:** 10.64898/2025.12.11.693746

**Authors:** Nataliia Kozhemiako, Naihua N. Gong, Shaun M. Purcell

## Abstract

Sleep electroencephalography (EEG) patterns exhibit complex variations influenced by multiple factors, including sex. While sex effects in sleep neurophysiology are relatively well described in adults, evidence in pediatric populations is more limited, particularly in the context of neurodevelopmental disorders (NDDs). Here, we analyzed a large pediatric clinical dataset to examine sex differences in whole-night sleep EEG among children without NDDs (N=1523, 2.5-17.5 years) and those with autism spectrum disorder (ASD, N=196), attention deficit hyperactivity disorder (ADHD, N=523), and intellectual disabilities (IntDis, N=167). Medical record analysis revealed limited sex differences in the prevalence of sleep disorders. Using the open-source package Luna, we extracted a broad range of sleep EEG features (435 in total) and found that 25% of all tested metrics showed significant sex differences in the non-NDD sample with the majority replicated in an independent population-based pediatric cohort. Non-NDD boys exhibited more fragmented sleep compared to girls, but this pattern was not observed among children with NDDs. In contrast, fast spindle density was lower in boys regardless of NDD diagnosis, which may contribute to males’ increased susceptibility to developing NDDs. No sex differences were observed in EEG-based brain age predictions. Several sleep EEG metrics showed nominal group-by-sex interactions with ASD and ADHD girls displaying more pronounced alterations, pointing to a varying degree of sleep disturbance. Overall, our findings demonstrate that many sleep EEG features exhibit significant sex differences during childhood, but these differences may be distinct in NDD populations. The sex-specific sleep alterations in NDDs underscore the importance of including both sexes in future research.

## Introduction

Sleep plays a crucial role in healthy childhood development (Blumberg et al., 2022). Although it remains unclear whether sleep problems are a cause or consequence of neurodevelopmental disorders (NDDs), they are notably more prevalent in these populations. In neurotypical children, the prevalence of sleep disorders is typically below 5% (Meltzer et al., 2010). In contrast, rates are significantly higher in children with NDDs – for example, 13% of children with autism spectrum disorder (ASD) (Lai et al., 2019) and 16% of children with mild to profound intellectual disabilities (Didden et al., 2002) exhibit sleep problems. Similarly, 25-50% of children with attention hyperactivity disorder (ADHD) experience sleep problems, with rates increasing with age (Cao et al., 2018; Wajszilber et al., 2018).

Another signature of NDDs is profound male preponderance (Zablotsky et al., 2019). Despite this phenomenon being well-documented, its neurophysiological underpinnings remain unclear. NDDs, including ASD, ADHD, intellectual disability (IntDis) have overlapping risk and prognostic factors, with male sex being a key one (Thapar et al., 2017). Increased male prevalence suggests the existence of early-life neurobiological factors that make boys vulnerable or, alternatively, are protective for girls. However, the specific genetic and hormonal contributions to these sex differences remain unclear (McCarthy et al., 2017). One prevailing hypothesis developed to explain the stark male preponderance in ASD, the female protective effect, posits that females require a higher burden of etiological risk factors to develop the disorder – a concept supported by genetic studies (Robinson et al., 2013; Wigdor et al., 2022). Sex differences also emerge in how NDDs present clinically, with males often exhibiting more severe symptoms and greater functional impairments (Allely, 2018; Arnett et al., 2015; Kittler et al., 2004; Loyer Carbonneau et al., 2021). However, it remains largely unclear whether sleep problems also exhibit sex-specific patterns in NDDs.

Sleep has been reported to differ between males and females including variations in the prevalence of sleep disorders, self-reported sleep patterns and sleep neurophysiology measured by polysomnography (Baker et al., 2020; Meers et al., 2019). While sex differences in sleep are well documented in adults (Chapman et al., 2025), findings in children are less consistent, likely because these differences are often assumed to arise mainly from hormonal changes during puberty (Mong & Cusmano, 2016). Therefore, studies examining sex differences in children’s sleep remain limited. Even without considering sex as a factor, research on sleep in NDDs faces notable challenges. A recent review highlighted likely roles of sleep oscillation abnormalities among NDDs but it also underscored 1) the scarcity of relevant studies and 2) limited sample sizes which, combined with 3) varying age ranges and 4) inconsistent methodologies, precluded strong conclusions (Gorgoni et al., 2020).

In this study, we examined how sex interacts with both PSG-derived sleep markers and clinically diagnosed sleep disorders in children with and without NDDs. Specifically, we focused on ASD, ADHD and IntDis, all of which are characterized by both male preponderance and increased prevalence of sleep problems. We used a large cohort of clinical pediatric sleep studies to comprehensively characterize sex differences in sleep across multiple levels: from sleep-related diagnoses to PSG-derived measures of sleep macro-architecture and microarchitecture including spectral characteristics of brain activity, sleep spindles, slow and infraslow oscillations. To better understand whether sleep alterations differ between boys and girls with NDDs, we compared each group separately to same-sex peers without NDDs and investigated sex-by-group interactions. Additionally, we applied a previously published model of brain age prediction in children to explore sex differences in brain age across groups and to identify whether brain age alterations in NDDs show sex-specific patterns relative to the non-NDD population.

## Methods

### Participants

In this study we used the data from two pediatric cohorts – the Nationwide Children’s Hospital Sleep DataBank (NCH) as a primary discovery dataset and the Child Adenotonsillectomy Trial (CHAT) as a replication dataset – both accessible through the National Sleep Research Resource (http://sleepdata.org). The NCH dataset was created to support research on pediatric sleep and includes individuals (N = 3,673) ranging from infancy to early adulthood who underwent clinical PSG at Nationwide Children’s Hospital between 2017 and 2019 (Lee et al., 2022). This dataset also contains demographic (age, sex, race), diagnostic information (ICD-9/10 codes) and medication records. All data were de-identified before being uploaded to the NSRR, and the dataset received exemption from the NCH Institutional Review Board along with a HIPAA waiver.

The CHAT dataset was created based on a multicenter study conducted across six pediatric clinical sites in the U.S., designed to screen children aged 5 to 9 years for a clinical trial targeting those who snored and were being evaluated for potential adenotonsillectomy. Participants were free of serious chronic health conditions and were not on ADHD medication (although 12 children had an ADHD diagnosis). All participants exhibited snoring and were considered possible candidates for adenotonsillectomy (Marcus et al., 2013; Weinstock et al., 2014). A total of 1,244 children underwent screening using PSG; for specifics on the inclusion criteria, see Marcus et al. (2013). For the current study, we analyzed data from the baseline PSG recordings only, excluding any follow-up studies conducted on children with mild to moderate obstructive sleep apnea. The CHAT study was approved by the respective Institutional Review Boards at each site, and written informed consent was obtained from each participant or their legal guardian.

We applied a set of primary exclusion criteria across both datasets. For the NCH sample, individuals were excluded if they were: (i) younger than 2.5 years (as many sleep EEG signatures are still emerging in that age range), (ii) older than 17.5 years (due to limited data in that age range), or (iii) diagnosed with narcolepsy. Within the non-NDD subsample, in addition to excluding individuals with ASD, ADHD, or IntDis, we also removed those with language disorders, tic disorders, cerebral palsy, Down syndrome, or epilepsy.

In the CHAT dataset, we excluded participants with missing age information and those diagnosed with ADHD. For both datasets, if an individual had multiple EEG recordings, only the first recording was retained for analysis.

### Diagnostic information

As outlined in our previous work using the NCH dataset (Kozhemiako et al., 2023), we extracted clinical information from the DIAGNOSIS.csv file. To identify specific diagnoses, we performed a string search using the diagnosis descriptions provided in the “DX_NAME” column. Detailed results of this search are presented in **Supplementary Table S1**. Additionally, we used data from the MEDICATION.csv file to account for medications prescribed at the time of the EEG recording. In particular, we controlled for medications known to affect sleep, including antihistamines (used by 114 girls and 176 boys), psychotropic medications (used by 109 girls and 142 boys), anticonvulsants (used by 77 girls and 96 boys), hormones (used by 77 girls and 99 boys), and sedatives/hypnotics (used by 1 girls and 5 boys, see **Supplementary Table S2**).

### Sleep EEG preprocessing

Whole-night sleep EEG recordings were processed using *Luna* (http://zzz.bwh.harvard.edu/luna/), an open-source software package developed by our team (S.M.P). We focused on six EEG channels – F3, F4, C3, C4, O1, and O2 – which were consistently available in both the NCH and CHAT datasets. For each recording, we selected 30-second epochs corresponding to specific sleep stages (N2, N3, and R) based on manual scoring aligned with AASM criteria.

Given the variation in original EEG sampling rates across and within datasets (256–512 Hz for NCH and 200–512 Hz for CHAT), all signals recorded above 200 Hz were down sampled to 200 Hz. Signals were re-referenced to the contralateral mastoids, converted to microvolt units, and bandpass filtered between 0.5 and 35 Hz. To address the significant line noise frequently observed in NCH recordings, we used a spectral interpolation method to remove interference, following the previously published approach (Leske & Dalal, 2019), as implemented in Luna.

As in Kozhemiako et al. (2023), we applied an automated procedure to exclude artifactual epochs. Within each sleep stage, epochs were removed if they exhibited maximum amplitudes above 200 µV or had flat or clipped signals for more than 10% of their duration. We also flagged epochs as outliers based on Hjorth parameters (activity, mobility, and complexity (Hjorth, 1970)): (i) any epoch exceeding 3 standard deviations from the individual’s channel-wide mean across all three parameters; (ii) values more than 4 SDs from the mean of other epochs within the same channel; or (iii) more than 4 SDs from the overall distribution across all channels. The Hjorth-based outlier detection was repeated twice per subject. Channels or epochs with over 50% of segments classified as outliers were excluded. These empirically determined thresholds aimed to eliminate prominent artifacts while retaining as much valid data as possible. Signal quality was further verified through visual inspection of randomly selected recordings and by examining spectral power distributions across frequencies for all signals.

### Sleep EEG final estimates

#### Spectral power

We computed spectral power using Welch’s method separately for each sleep stage (N2, N3, and R), summarizing the results within standard frequency bands: slow (0.5–1 Hz), delta (1–4 Hz), theta (4–8 Hz), alpha (8–12 Hz), sigma (12–15 Hz), beta (15–30 Hz). For each 30-second epoch, we applied a Fast Fourier Transform (FFT) using 4-second segments, yielding a spectral resolution of 0.25 Hz. A 50% Tukey taper was used to window each segment, and segments overlapped by 50% (2 seconds). Power was first averaged across segments within each epoch, then averaged again across all epochs within a given channel and sleep stage. Relative power was calculated as the proportion of power in each frequency band relative to the total absolute power. Absolute power values were log-transformed before statistical analysis.

#### Spindles

As in Kozhemiako et al. (2023), we detected the two spindle types separately given a recent evidence that slow frontal and fast central spindles can be distinguished as early as 18 months of age (Kwon et al., 2022). Detection was performed using 7-cycle complex Morlet wavelets centered at 11 Hz (slow spindles or SS) and 15 Hz (fast spindles or FS), each spanning approximately ±2 Hz, following previously published methods (Purcell et al., 2017; Warby et al., 2014).

Putative spindles were identified from temporally smoothed (0.1 s window) wavelet coefficients using two criteria: intervals had to exceed 4.5× the mean amplitude for at least 300 ms and 2× the mean for at least 500 ms. Segments longer than 3 seconds were discarded, and intervals within 500 ms were merged unless the merged segment exceeded the 3-second limit.

To improve specificity, we applied a quality control step that excluded spindles where the increase in non-spindle band power (delta, theta, beta) exceeded the increase in spindle-band (sigma) power, relative to N2 sleep. This ensured that retained events reflected true spindle activity rather than general signal amplitude increases, which may indicate artifacts.

From the spindles that passed QC, we computed spindle density (count per minute), amplitude, duration, peak frequency, and chirp—the change in frequency from the first to the second half of the spindle, with negative values indicating deceleration.

#### Slow oscillations

Slow oscillations (SOs) were detected from EEG signals filtered between 0.5 and 4 Hz by identifying zero-crossings. To qualify as a putative SO, events had to meet two temporal criteria: (1) the duration between zero-crossings surrounding the negative peak had to be between 0.3 and 1.5 seconds, and (2) the segment leading to the positive peak could not exceed 1 second. Then we used a relative threshold requiring both the negative peak and the peak-to-peak amplitude to exceed twice the individual/channel-specific mean. For each channel, we calculated SO density (events per minute), along with average negative peak amplitude, peak-to-peak amplitude, event duration, and the slope of the rising phase from the negative peak.

#### Spindles and slow oscillation coupling

Based on detected spindles and SOs, we quantified their coupling for each EEG channel using three metrics. First, we calculated the proportion of spindles that coincided with an SO, referred to as “gross overlap.” Next, we used the filter-Hilbert method to estimate the SO phase at the spindle peak and computed the average phase across events for each channel (i.e., the coupling phase angle). Lastly, we assessed the consistency of phase alignment between SOs and spindles using inter-trial phase clustering, which reflects the strength of phase coupling (coupling magnitude).

To assess statistical significance, the gross overlap and coupling magnitude values were z-transformed using null distributions generated from 10,000 random permutations. In each permutation, the time indices were shuffled while preserving the number of SOs, spindles, and (for coupling magnitude) their overall overlap.

#### Infraslow oscillations (ISO)

Continuous N2 sleep bouts (≥256 s) were divided into overlapping 4-s epochs (25% overlap). For each, power spectral density was computed after linear detrending and Hanning tapering, then averaged across seven bands: slow (0.5–1 Hz), delta (1–4 Hz), theta (4–7 Hz), alpha (7–10 Hz), slow sigma (10–12 Hz), fast sigma (12–15 Hz), and beta (15–30 Hz). This produced infraslow time series for each band (1-s resolution). Each series was then bandpass filtered (0.01–0.03 Hz, FIR filter; 1% ripple, 0.005-Hz transition) to capture the infraslow rhythm, centered near 0.02 Hz as reported previously (Lecci et al., 2017). Each band’s time series underwent multitaper spectral analysis with spectral densities normalized to total power in the 0–0.1 Hz range, and infraslow power computed as the length-weighted average of these relative spectral densities across all N2 bouts. We estimated amplitude and frequency of the ISO spectral power peak for each frequency band.

#### Predicted Age based on sleep EEG

We predicted age based on sleep EEG features using similar approach as in *Kozhemiako et al. 2023.* We trained a multiple linear regression model using the sleep metrics analyzed in this study. After excluding individuals from NDD subgroups, the non-NDD NCH sample was randomly split into a training set (70%) and a held-out test set (30%). To reduce feature redundancy, we removed highly correlated variables (absolute r > 0.9) based on the training data. The remaining features were z-scored across all datasets using the mean and standard deviation from the training set. Because our goal was to explore sex-related effects, sex was not included as a covariate in the model, though race was. The model was evaluated on four independent test sets: the non-NDD NCH held-out sample, and the ASD, ADHD, and IntDis subgroups. Performance was assessed using Pearson’s correlation, mean absolute error (MAE), and mean prediction error between predicted and actual chronological age.

#### Final exclusion criteria based on sleep EEG

For both datasets, we applied a set of final exclusion criteria to ensure sufficient data quality based on EEG features. First, recordings with total sleep time (TST) under 180 minutes were excluded. Further exclusions were made based on the following criteria: (i) fewer than 10 valid epochs remaining in any sleep stage (N2, N3, R) after artifact rejection, (ii) persistent line noise (defined as SPK > 5 SD in any channel during any stage), and (iii) extreme spectral power values – specifically, power at 1 Hz falling outside ±4 SD in any channel (to capture movement, eye artifacts, or poor signal quality) or power at 25 Hz exceeding +4 SD (to detect muscle activity artifacts).

Signal polarity inconsistencies were noted in some recordings, requiring additional filtering for analyses sensitive to polarity, such as those involving SOs and their coupling with spindles. Recordings with ambiguous polarity (T_DIFF between –1 and 1 at C3 or C4 during N2, per Luna’s POL command) were excluded. For recordings where T_DIFF exceeded 1 at either site, polarity was flipped to ensure consistency.

Final sample sizes and demographic details for the analyzed cohorts are provided in Table 1 and Supplementary Table S3.

**Table 1.**
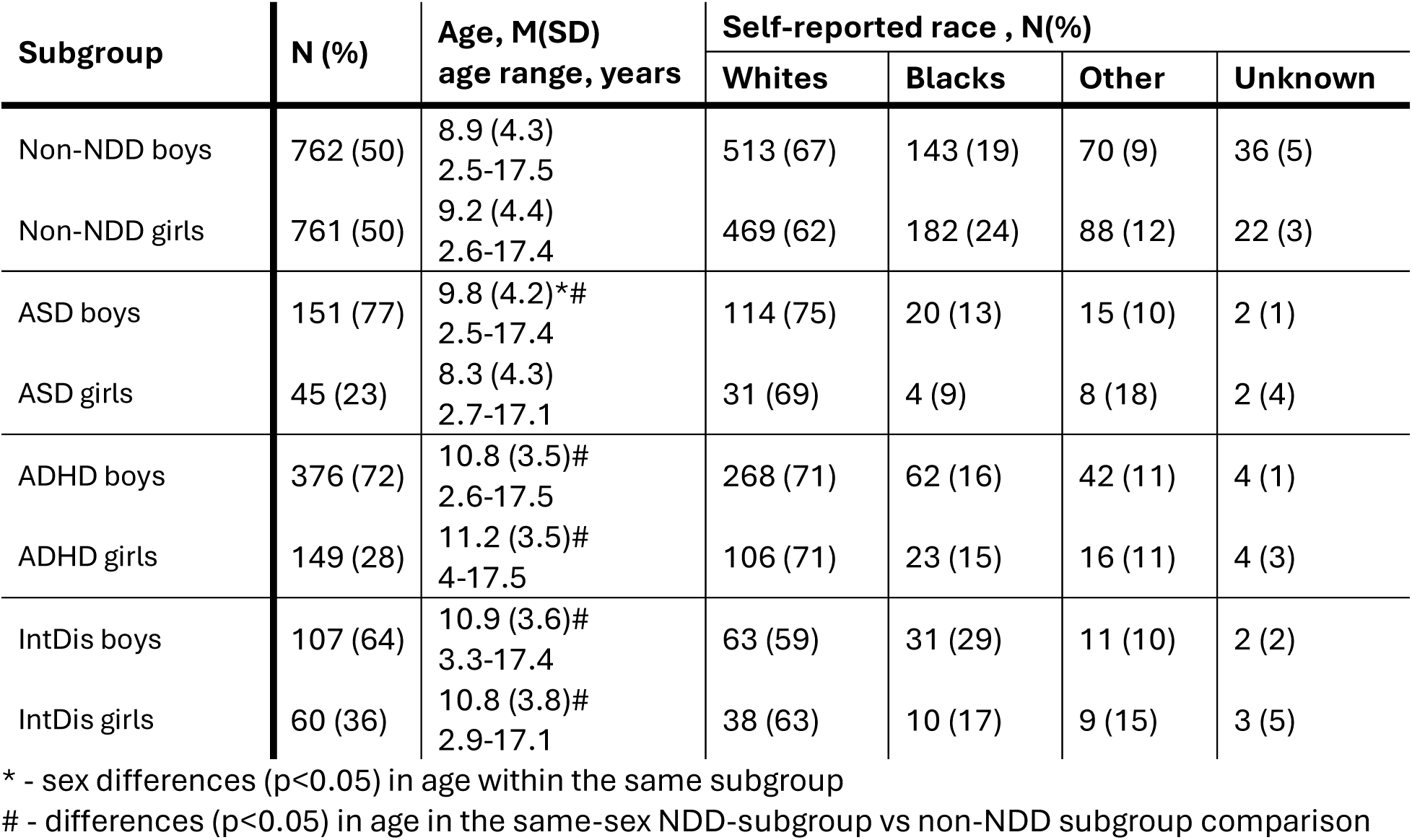
Demographic characteristics of the NCH sample.

| Subgroup | N (%) | Age, M(SD)<br>age range, years | Self-reported race , N(%) |  |  |  |
| --- | --- | --- | --- | --- | --- | --- |
|  |  |  | Whites | Blacks | Other | Unknown |
| Non-NDD boys | 762 (50) | 8.9 (4.3)<br>2.5-17.5 | 513 (67) | 143 (19) | 70 (9) | 36 (5) |
| Non-NDD girls | 761 (50) | 9.2 (4.4)<br>2.6-17.4 | 469 (62) | 182 (24) | 88 (12) | 22 (3) |
| ASD boys | 151 (77) | 9.8 (4.2)*#<br>2.5-17.4 | 114 (75) | 20 (13) | 15 (10) | 2 (1) |
| ASD girls | 45 (23) | 8.3 (4.3)<br>2.7-17.1 | 31 (69) | 4 (9) | 8 (18) | 2 (4) |
| ADHD boys | 376 (72) | 10.8 (3.5)#<br>2.6-17.5 | 268 (71) | 62 (16) | 42 (11) | 4 (1) |
| ADHD girls | 149 (28) | 11.2 (3.5)#<br>4-17.5 | 106 (71) | 23 (15) | 16 (11) | 4 (3) |
| IntDis boys | 107 (64) | 10.9 (3.6)#<br>3.3-17.4 | 63 (59) | 31 (29) | 11 (10) | 2 (2) |
| IntDis girls | 60 (36) | 10.8 (3.8)#<br>2.9-17.1 | 38 (63) | 10 (17) | 9 (15) | 3 (5) |

\* - sex differences ( $p < 0.05$ ) in age within the same subgroup

### Statistical analysis

To assess sex differences in each sleep EEG metric, we used linear regression models in the non-NDD NCH and CHAT samples. Sleep EEG features served as dependent variables, with sex as the main predictor. Age and self-reported race were included as covariates. For the NCH sample, we also controlled for sleep disorder diagnoses and medication use. The latter was modelled as a binary variable of prescribed medication coinciding with sleep EEG and included anticonvulsants, antihistamines, psychotherapeutics, sedatives and melatonin. Prior to modeling, we removed outliers greater than 3 standard deviations from the mean. Sex differences within the ASD, ADHD, and IntDis groups were examined using a similar model, with additional adjustment for overlapping NDD disorder diagnoses (e.g., children with concurrent ASD, ADHD and/or IntDis diagnoses).

To examine sex differences in the prevalence of sleep disorder diagnoses in the NCH sample, we employed logistic regression, treating diagnosis as the binary dependent variable and sex as the predictor, adjusting for age, self-reported race, medication use, and disorder overlap (for NDD groups).

We also evaluated diagnostic group effects (NDD vs. non-NDD) separately for boys and girls using the same modeling approach, replacing sex with diagnosis as the binary predictor while keeping all covariates. In addition, we tested for sex-by-group interactions for each EEG metric.

Given the large number of tests, we corrected p-values using the False Discovery Rate (FDR) procedure (Benjamini & Hochberg, 1995) with an alpha level of 0.1. Corrections were applied separately for three sets of analyses: (1) sex effects (351 EEG features × 4 groups), (2) group effects within sex (351 features × 3 group comparisons), and (3) sex-by-group interactions (351 features × 3 group comparisons).

## Results

The final sample included four subgroups divided by sex (**Table 1**). The non-NDD subgroup was balanced by sex, whereas the NDD groups – as expected – had a higher proportion of boys. Although the age range was broadly similar across all groups (3 – 17 years), children with NDDs were on average older than their same-sex non-NDD peers. Additionally, within the ASD group, girls were younger than boys. To account for these differences, we included age and other relevant covariates in all subsequent analyses.

### Limited sex differences in sleep disorder prevalence within groups, but markedly elevated rates in both boys and girls with NDDs compared to non-NDD group

After adjusting for age, race, and medication use (and overlap in NDD diagnoses in the NDD subgroups comparison), modest sex differences in the rates of sleep disorders were observed only in two groups (**Figure 1**). In the ADHD group, boys had higher rates of sleep apnea compared to girls (b = 0.42, *p* = 0.044). In the non-NDD group, girls had higher rates of "other sleep problems" – a broad category that includes unspecified sleep disorder diagnoses, narcolepsy, night terrors, and others (see Table S1) – compared to boys (b = 0.34, *p* = 0.007). Given reports suggesting that sex difference in sleep disorders appear during puberty, we repeated the analysis in a subsample of non-NDD individuals older than 12 years (231 girls, 197 boys) and found that in addition to higher rate of various sleep disorders under umbrella category “other disorders”, girls also had higher rate of circadian disorder (b = 0.64, *p* = 0.025) while boys had higher rate of apnea (b = 0.71, *p* = 0.002).

**Figure 1.**
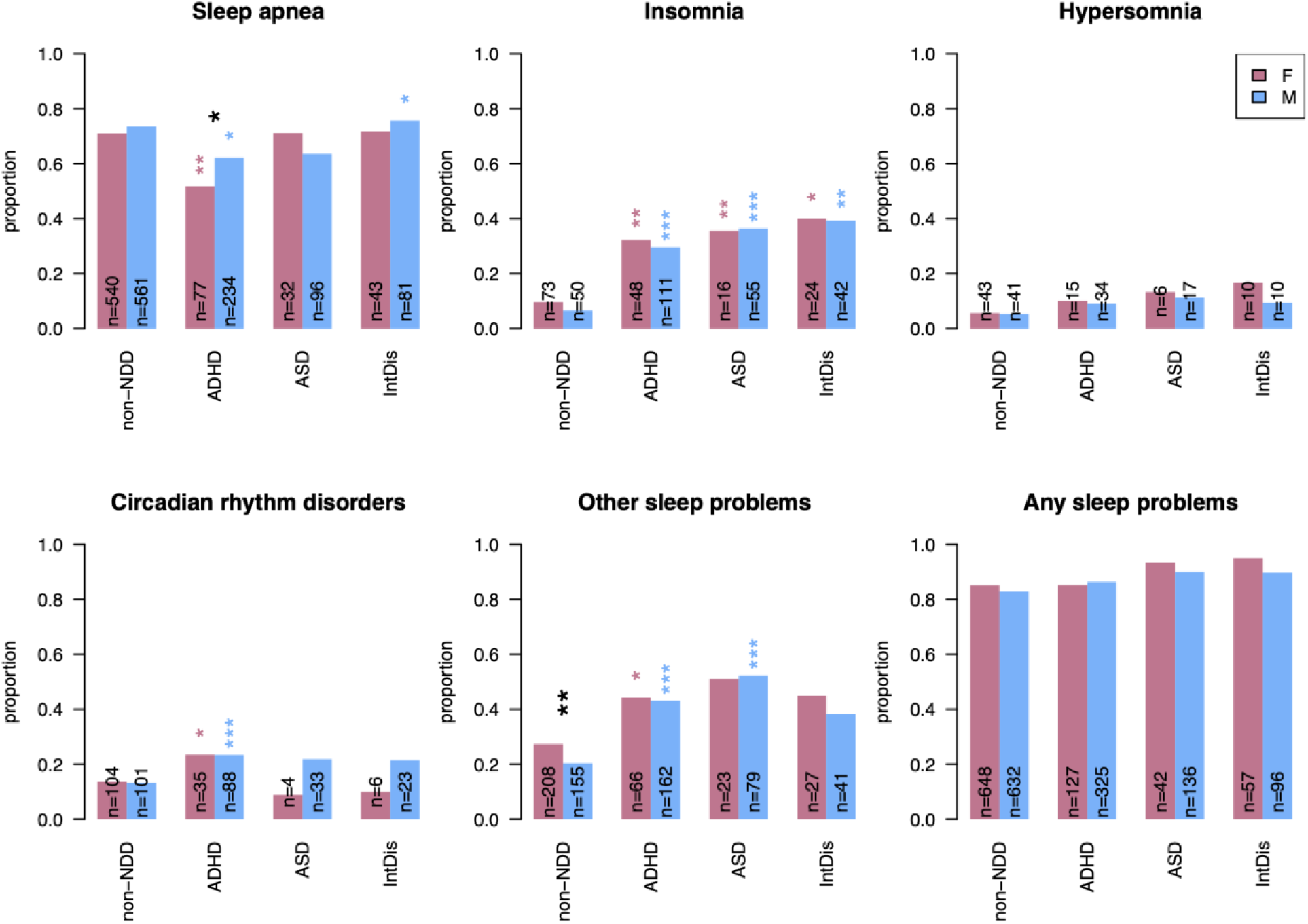
Differences in sleep disorders in the NCH cohort. The bar plots illustrate the proportion of individuals with sleep diagnoses separately in boys and girls. Significant sex differences after controlling for age, race and medication within a diagnostic group are marked with black stars, while same-sex comparison between a NDD subgroup and non-NDD individuals is shown with pink (for girls) or blue (for boys) stars. * – p < 0.05; ** – p < 0.01; *** – p < 0.001

When comparing NDD and non-NDD groups within the same sex, both boys and girls with NDDs had higher rates of insomnia than their non-NDD counterparts (b = 0.77 – 1.57, all p < 0.05). Additionally, both boys and girls with ADHD showed significantly higher rates of circadian rhythm disorders (b=0.66, p = 0.0008 in boys, b=0.52, p = 0.04 in girls) and other sleep problems (b=0.84, p = 5x10^-7^ in boys, b=0.42, p = 0.042 in girls), but lower rates of sleep apnea (b=-0.35 p = 0.03 in boys, b=-0.7, p = 0.001 in girls). In the ASD group, both boys and girls had higher rates of other sleep problems (b=1.14 p = 5x10^-5^ in boys, b=0.38, p = 0.041 in girls), while boys with IntDis had higher rates of sleep apnea (b=0.99, p = 0.035) compared to the non-NDD boys. However, when explicitly testing for group-by-sex interactions, no significant interaction effects were observed for any diagnostic group.

### Numerous sleep EEG metrics differ significantly between non-NDD boys and girls

After multiple comparison correction (MCC), 25% of all tested metrics showed significant sex differences between boys and girls in the non-NDD sample (Figure 2A). The largest effect sizes were observed in absolute alpha and sigma power in occipital channels during R sleep, which were higher in girls (maximum effect size at O1: *b* = –0.25, *p* = 6 × 10⁻⁷; **Figure 2B**).

**Figure 2.**
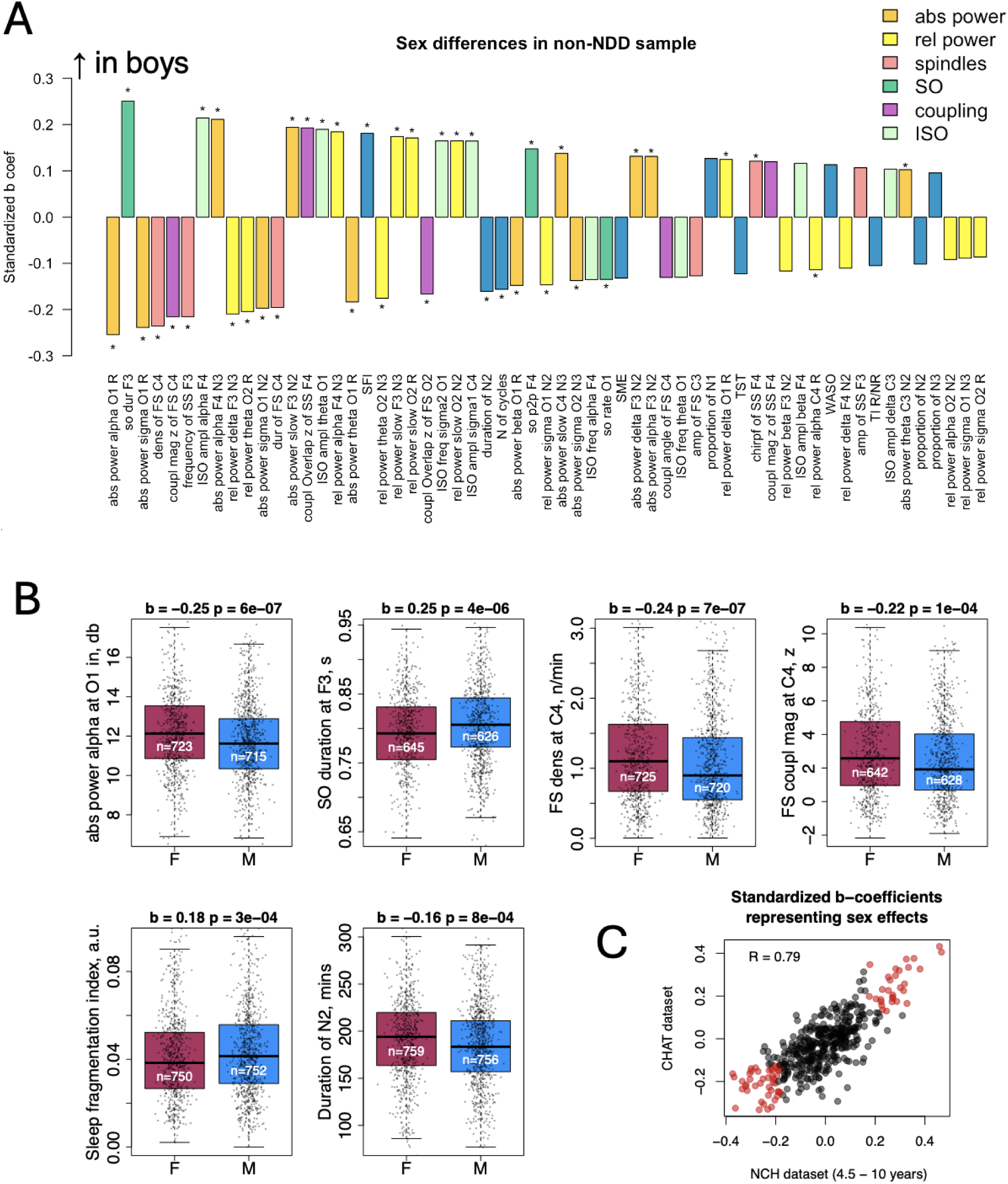
Sex differences in sleep EEG metrics in the non-NDD sample. A – The bar plot illustrates all sleep EEG metrics with nominally significant sex differences (p < 0.05, star marks comparisons with significant sex differences after MCC. The metrics are sorted by their absolute standardized b-coefficient representing the effect size. B – the box plots provide examples of the sleep EEG metrics with the largest effect sizes. C – the scatterplot showing replication of sex differences in the CHAT dataset. Red dots indicate sleep metrics that showed significant sex differences (after multiple comparisons correction) in the age-matched non-NDD subsample of the NCH dataset and were also nominally significant in the CHAT dataset.

In contrast, alpha power in frontal channels during NR sleep was higher in boys, with the strongest effect at F4 during N3 (*b* = 0.21, *p* = 3 × 10⁻⁵). Other prominent differences included longer SO duration in boys (maximum effect at F3: *b* = 0.25, *p* = 4 × 10⁻⁶), lower FS density (*b* = –0.24, *p* = 7 × 10⁻^7^ at C4), and reduced coupling magnitude between FS and SOs (*d* = –0.22, *p* = 1 × 10⁻^4^ at C4; Figure 2B).

In terms of sleep macroarchitecture, boys generally exhibited poorer sleep quality, as indicated by higher sleep fragmentation index (SFI, *b* = 0.18, *p* = 3 × 10⁻^4^), shorter total N2 stage duration (*b* = −0.16, *p* = 8 × 10⁻^4^) and fewer sleep cycles (*b* = −0.16, *p* = 9 × 10⁻^4^).

We used the Child Adenotonsillectomy Trial (CHAT, N= 1,201, 630 females) as a replication dataset to verify if sex differences expressed in the non-NDD group were consistent. Since the CHAT dataset has a narrower age range (4.5 – 10 years, mean age 7.1 years), we selected non-NDD NCH individuals of similar age (N = 1161, 499 females, 4.5 – 10 years, mean age 7.1 years), and compared sex differences in both samples. Overall, 75 out of 93 sleep EEG metrics significant after MCC in this subset of the NCH cohort were also nominally significant (p < 0.05) in the CHAT dataset and the sex differences were in the same direction (**Figure 2C**). Additionally, the standardized b-coefficients capturing sex effects in both datasets were highly correlated (R = 0.79) suggesting sex differences are highly consistent across cohorts.

Of note, the number of sleep EEG metrics with significant sex differences after MCC in this prepubertal (< 10 years) subset of the NCH cohort was comparable to the whole sample (93 vs 107, with 63 metrics being significant in both) contrary to the common belief that sex differences arise primarily after puberty.

### Sex differences in NDDs only partially overlap with those in non-NDD groups

In NDD groups, smaller number of metrics expressed nominally significant (p < 0.05) differences between boys and girls (11% in ADHD, 7% in ASD and 4% in IntDis) compared to non-NDD individuals (41%), however, this may be explained by lower statistical power to detect those differences due to smaller sample size in NDDs. Most sex differences in the ADHD and ASD groups overlapped with those seen in the non-NDD group, whereas in the IntDis group, the proportion of overlapping and unique sex differences was approximately equal (**Figure 3A**).

**Figure 3.**
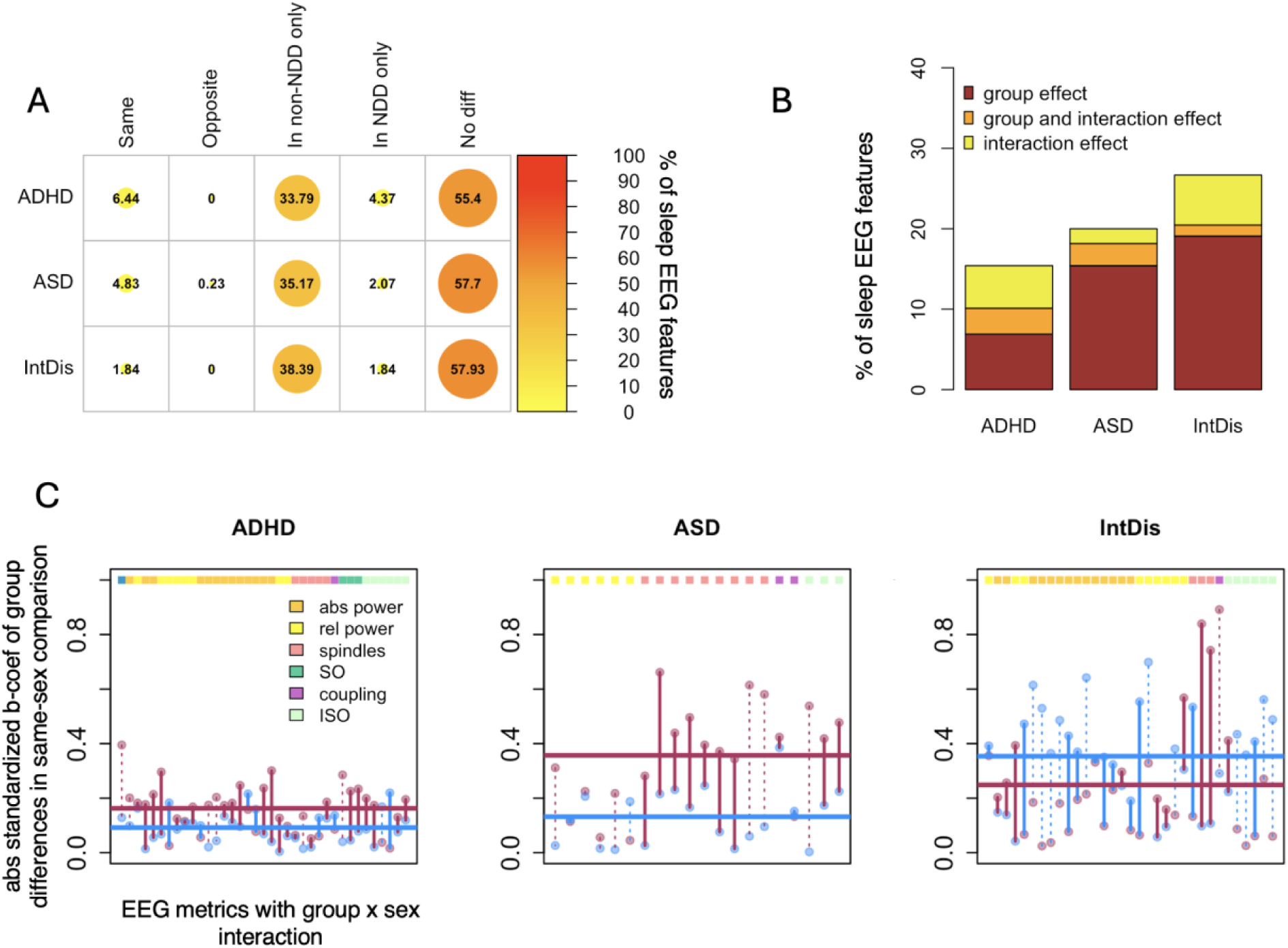
Sex differences in NDDs only partially overlap with those in non-NDD groups and case-control differences express sex-specific effects. A – The plot displays the percentage of sleep EEG metrics with nominally significant sex differences (p < 0.05), categorized as present in both a specific neurodevelopmental disorder (NDD) group and the non-NDD group with effects in the same or opposite directions, present only in the non-NDD group, present only in the specific NDD group, or absent in both groups. B – The box plots show the percentage of sleep EEG metrics with nominally significant (p < 0.05) effects for group differences between a specific NDD group and the non-NDD group, group-by-sex interaction effects, and overlapping effects of group differences and group-by-sex interactions. C – The scatterplot compares sex-specific effect sizes of NDD-related alterations, measured as absolute standardized β-coefficients from linear regression models testing group differences in same-sex comparisons between a specific NDD group and the non-NDD group, for ADHD, ASD, and Intellectual Disability (IntDis) groups. Horizontal lines indicate the average effect size of group differences for males (blue) and females (maroon). Vertical lines show whether group differences are in the same (dotted) or opposite (solid) directions, with colors highlighting whether the absolute effect size is larger in females (maroon) or males (blue).

When examining the effects of NDDs on sleep EEG, we found that individuals with IntDis showed the highest proportion of altered sleep features compared to the non-NDD group (21% of EEG metrics with nominally significant differences; **Figure 3B**). The proportions were 17% in the ASD group and 12% in the ADHD group.

For sex-by-group interactions, the ADHD group showed the largest proportion of nominally significant metrics across domains (9%), followed by IntDis (8%, mainly spectral power metrics) and ASD (5%, mainly relative power and spindle metrics).

We observed stronger sleep alterations in ASD and ADHD girls compared to boys within the same-sex comparisons to non-NDD children (**Figure 3C**), which may be consistent with the female protective effect theory. However, the pattern was the opposite in children with IntDis, where boys showed larger effect sizes than girls suggesting that sex-specific alterations may vary across different NDDs.

### Sex differences in sleep EEG metrics in ADHD

In the ADHD subsample, sex differences were observed across multiple sleep metrics (**Figure 4 A**), partially mirroring patterns seen in the non-NDD group. For example, similar to the non-NDD group, boys with ADHD showed lower FS density (largest effect at C4: *b* = −0.45, *p* = 2 × 10⁻⁵) and shorter FS duration (largest effect at O1: *b* = –0.29, *p* = 0.005) across several channels. They also exhibited lower relative sigma power during the N2 stage (at C3: *b* = - 0.28, *p* = 0.004).

**Figure 4.**
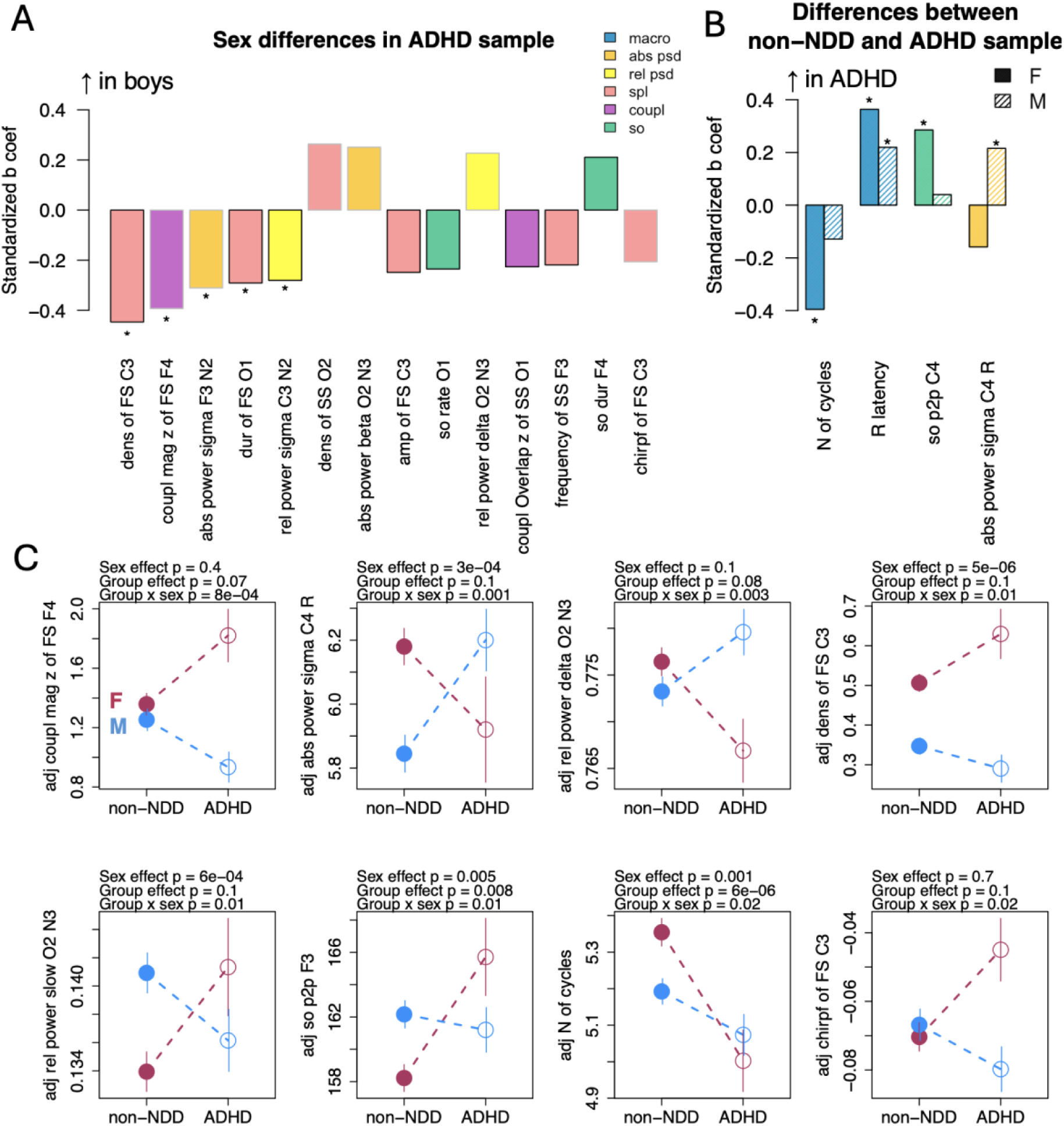
Sex differences in sleep EEG metrics in the ADHD sample. A – Bar plot showing standardized b-coefficients for all nominally significant sex differences between boys and girls with ADHD, ordered by absolute effect size. For metrics recorded across multiple channels, only the channel with the largest effect size is displayed. A star indicates metrics with sex differences that remained significant after MCC. Bar outline color reflects whether a similar sex difference was observed in the non-NDD group: black outlines indicate the same direction of sex difference as in the non-NDD group; grey outlines indicate that no sex difference was observed in the non-NDD group for that metric; B – Bar plot showing standardized b-coefficients for all metrics with significant group differences (after MCC) in ADHD relative to same-sex non-NDD peers. Solid bars indicate effects in girls; dashed bars indicate effects in boys. Stars indicate whether significant group differences were found in girls, boys, or both; C – Examples of nominally significant group-by-sex interactions.

In contrast to the non-NDD group, where significant sex differences in absolute sigma power were primarily found in central and occipital channels, the ADHD group showed differences in frontal regions (at F3: *b* = –0.31, *p* = 0.006). A similar topographical distinction was observed in FS–SO coupling magnitude: in ADHD, lower coupling in boys was found in frontal channels (at F4: *b* = –0.39, *p* = 3 × 10⁻⁴), whereas in the non-NDD group, this effect was localized to central regions. These distinct sex effects in ADHD vs non-NDD groups were also observed as nominally significant group-by-sex interactions (**Figure 4 C**).

The comparison between ADHD and non-NDD participants within the same sex revealed significant differences in four sleep EEG metrics (**Figure 3 B**). Stage R latency was increased in both girls and boys with ADHD compared to their non-NDD counterparts (*b* = 0.26, *p* = 4 × 10⁻⁴ for girls; *b* = 0.22, *p* = 0.004 for boys). Sex-specific differences included fewer sleep cycles in girls with ADHD (*b* = –0.4, *p* = 5 × 10⁻⁵) and higher SO peak-to-peak amplitude (*b* = 0.29, *p* = 0.001) compared to non-NDD girls, two effects that were not seen in boys. Conversely, in boys with ADHD, absolute sigma power during R sleep was reduced relative to non-NDD boys (*b* = 0.22, *p* = 0.007), which was not observed in girls. All these three metrics expressed nominally significant group-by-sex interactions (**Figure 4 C**).

### Sex differences in sleep EEG metrics in ASD

In the ASD subsample, the only metric showing a significant sex difference after MCC was the shorter SS duration in boys compared to girls, with the largest effect at O2 (*b* = –0.56, *p* = 0.004), despite many other metrics showing nominal significance (**Figure 5A**). Notably, this sex difference in SS duration was not observed in the non-NDD group. Another interesting finding was the opposite pattern of sex differences in SS frequency. In the ASD group, boys showed nominally higher FS frequency at C3 (*b* = 0.41, *p* = 0.02), whereas in the non-NDD group, boys showed lower FS frequency (*b* = –0.13, *p* = 0.006). Such sex specific effect in SS duration and frequency were confirmed by nominally significant group-by-sex interaction terms (**Figure 5C**).

**Figure 5.**
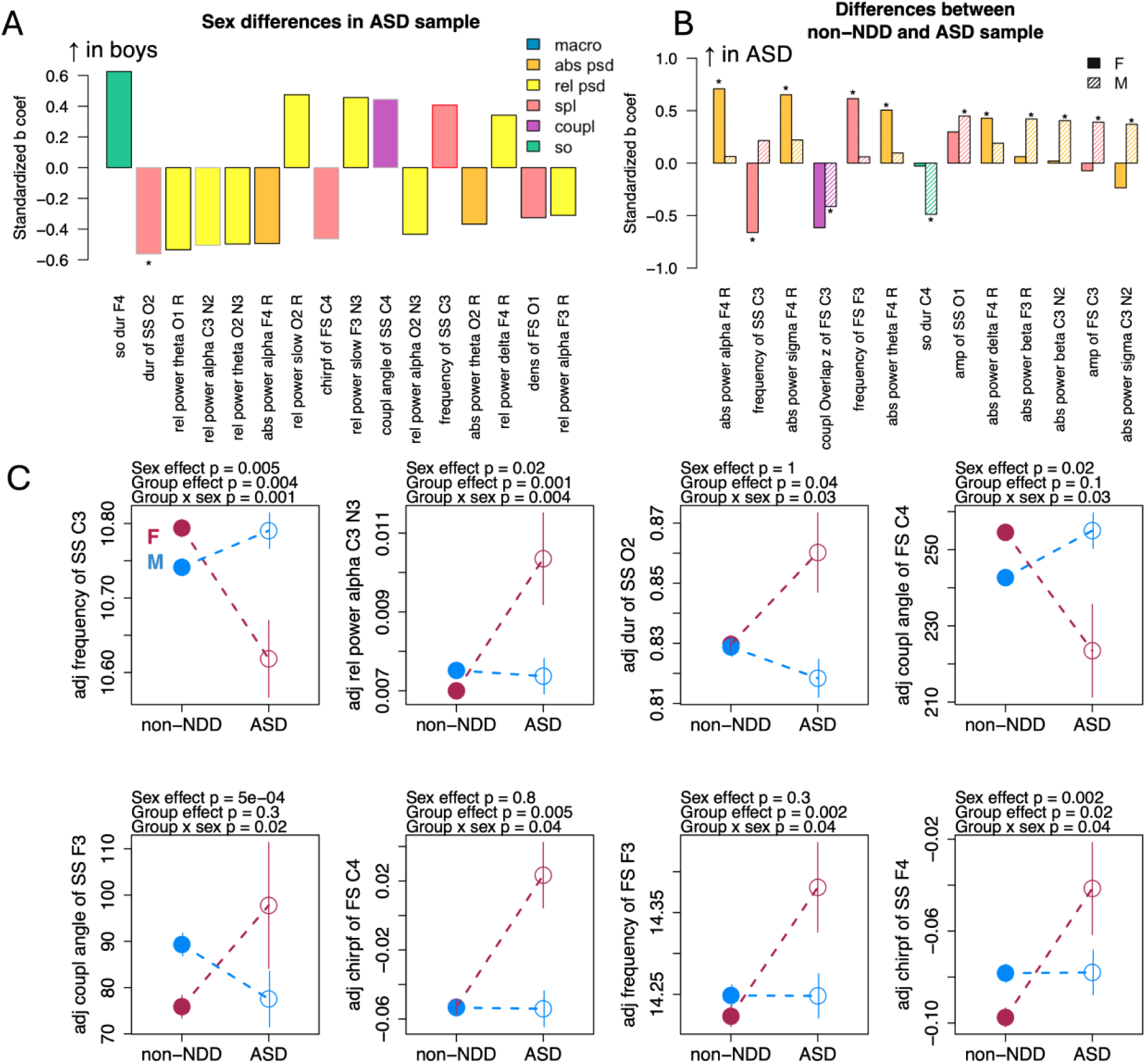
Sex differences in sleep EEG metrics in the ASD sample. A – Bar plot showing standardized b-coefficients for all nominally significant sex differences between boys and girls with ASD, ordered by absolute effect size. For metrics recorded across multiple channels, only the channel with the largest effect size is displayed. A star indicates metrics with sex differences that remained significant after MCC. Bar outline color reflects whether a similar sex difference was observed in the non-NDD group: black outlines indicate the same direction of sex difference as in the non-NDD group; grey outlines indicate that no sex difference was observed in the non-NDD group for that metric; B – Bar plot showing standardized b-coefficients for all metrics with significant group differences (after MCC) in ASD relative to same-sex non-NDD peers. Solid bars indicate effects in girls; dashed bars indicate effects in boys. Stars indicate whether significant group differences were found in girls, boys, or both; C – Examples of nominally significant group-by-sex interactions.

Same-sex comparisons between the ASD and non-NDD groups revealed multiple significant differences, most of which were specific to sex (**Figure 5B**). The only metric showing a consistent pattern across sexes was the reduced coupling overlap between SO and FS at the C3 channel, which was significantly lower in boys (*b* = –0.41, *p* = 0.008) and nominally reduced in girls (*b* = –0.61, *p* = 0.01).

Girls with ASD showed a broad increase in absolute power across multiple frequency bands during R, suggesting a general elevation in total power compared to non-NDD girls, with the largest effect in the alpha band at F4 (*b* = 0.71, *p* = 5 × 10⁻⁴). They also exhibited lower SS (C3: *b* = –0.66, *p* = 0.001) but higher FS frequency (F3: *b* = 0.61, *p* = 0.002), particularly in frontal and central regions while ASD boys were not different from their non-NDD peers. Both SS and FS frequency exhibited nominally significant group-by-sex interaction (**Figure 5C**).

Boys with ASD, on the other hand, displayed elevated power primarily in higher frequency bands, with the strongest effect in beta power during R sleep at F3 (*b* = 0.42, *p* = 0.001, **Figure 5B**). This increase in power at higher frequencies was also reflected in greater spindle amplitude, notably for slow spindles at O1 (*b* = 0.45, *p* = 7 × 10⁻⁴). Boys with ASD also had shorter SO duration compared to non-NDD boys (C4: *b* = –0.49, *p* = 0.002).

### Sex differences in sleep EEG metrics in IntDis

In the IntDis group, boys and girls showed only nominally significant sex differences, none of which survived MCC. Some of these sex differences, such as reduced FS density in boys, were also observed in the non-NDD, ADHD, and ASD groups (**Figure 6A**).

**Figure 6.**
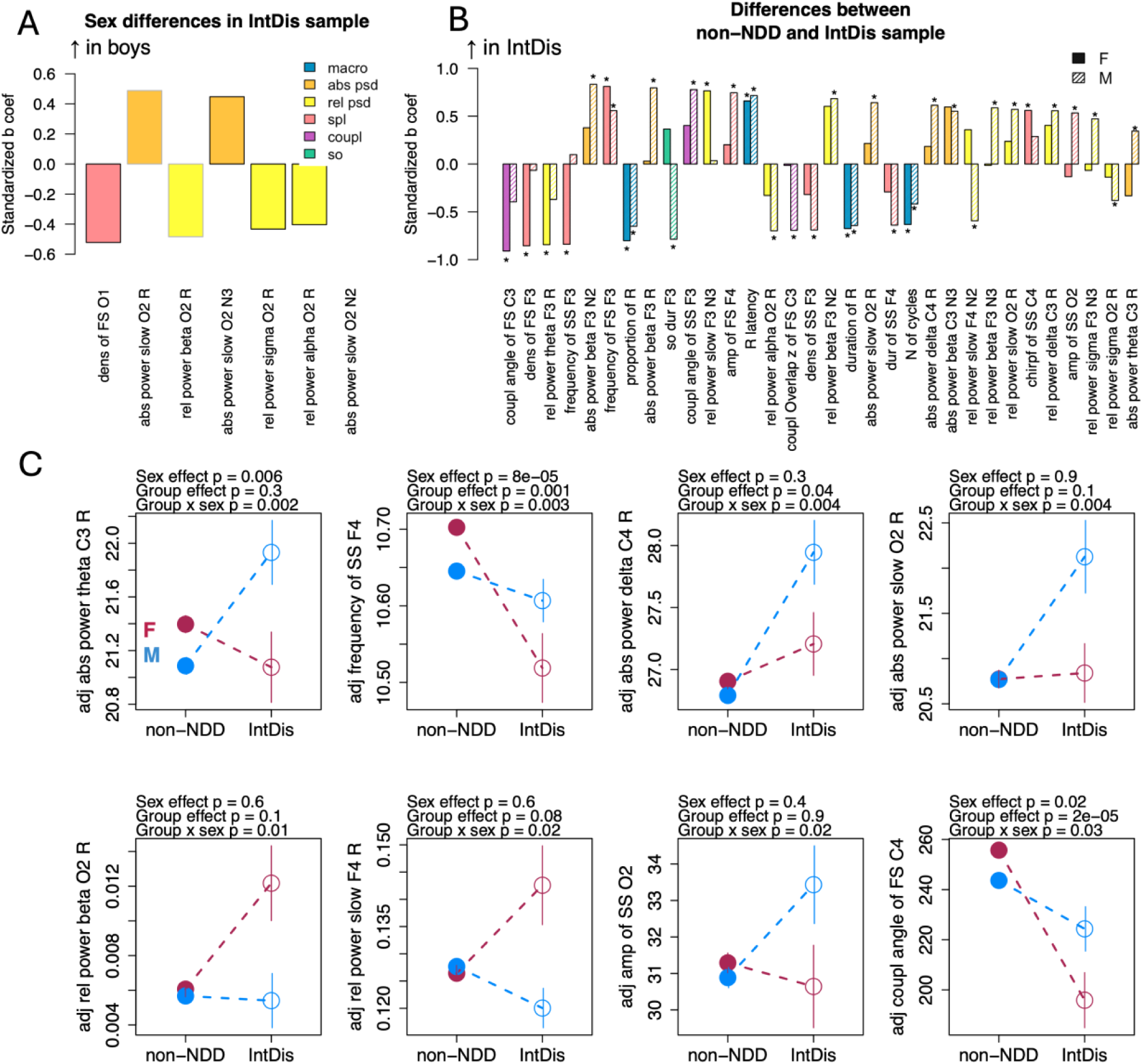
Sex differences in sleep EEG metrics in the IntDis sample. A – Bar plot showing standardized b-coefficients for all nominally significant sex differences between boys and girls with IntDis, ordered by absolute effect size. For metrics recorded across multiple channels, only the channel with the largest effect size is displayed. A star indicates metrics with sex differences that remained significant after MCC. Bar outline color reflects whether a similar sex difference was observed in the non-NDD group: black outlines indicate the same direction of sex difference as in the non-NDD group; grey outlines indicate that no sex difference was observed in the non-NDD group for that metric; B – Bar plot showing standardized b-coefficients for all metrics with significant group differences (after MCC) in IntDis relative to same-sex non-NDD peers. Solid bars indicate effects in girls; dashed bars indicate effects in boys. Stars indicate whether significant group differences were found in girls, boys, or both; C – Examples of nominally significant group-by-sex interactions.

Despite the lack of strong sex differences within the IntDis group, both boys and girls showed the highest number of group differences when compared to their same-sex peers in the non-NDD group (**Figure 6B**). Sleep macroarchitecture was similarly altered in both sexes: individuals with IntDis had shorter R sleep (as a proportion of total sleep time [TST]; girls: *b* = –0.80, *p* = 4 × 10⁻⁶; boys: *b* = –0.65, *p* = 0.004), longer R latency (girls: *b* = 0.66, *p* = 6 × 10⁻⁴; boys: *b* = 0.72, *p* = 2 × 10⁻⁵), and fewer sleep cycles (girls: *b* = –0.63, *p* = 5 × 10⁻⁴; boys: *b* = – 0.42, *p* = 0.005). In addition, both boys and girls with IntDis had significantly elevated FS frequency across all channels, with the largest effects at F3 (girls: *b* = 0.81, *p* = 3 × 10⁻⁵; boys: *b* = 0.56, *p* = 9 × 10⁻⁴). In contrast, SS frequency was significantly reduced compared to non-NDD, but only in girls with IntDis (F3: *b* = –0.84, *p* = 1 × 10⁻⁵) presenting a similar sex specific pattern to ASD group also confirmed by nominally significant group-by-sex interaction term (**Figure 6C**).

Several other sleep metrics were significantly altered in either boys or girls with IntDis, but not both. For example, girls with IntDis showed reduced FS density (F3: *b* = –0.86, *p* = 4 × 10⁻⁵), whereas boys had reduced SS density (F3: *b* = –0.69, *p* = 2 × 10⁻⁴) compared to same-sex non-NDD participants. Another example included coupling characteristics between spindles and SO: in girls, FS occurred earlier in the SO cycle in central regions (C3: *b* = –0.91, *p* = 2 × 10⁻⁴), while in boys, SS overlapped with SO at a later phase (F3: *b* = 0.78, *p* = 2 × 10⁻⁵) compared to their non-NDD peers. However, there was no significant group-by-sex interactions detected for these metrics except FS coupling angle (**Figure 6C**).

Additional findings in boys with IntDis included increased absolute beta power across all channels and stages (maximum effect at F3 during N2: *b* = 0.83, *p* = 1 × 10⁻⁶), higher slow and delta power during R (O2: *b* = 0.64, *p* = 9 × 10⁻⁶, also showing nominally significant group-by-sex interactions), greater SS and FS amplitude (maximum effect for FS at F4: *b* = 0.75, *p* = 6 × 10⁻⁴), and shorter SO duration in frontal and central channels (F3: *b* = –0.75, *p* = 2 × 10⁻⁴) compared to non-NDD boys.

In contrast, girls with IntDis exhibited lower relative theta power during R (F3: *b* = –0.84, *p* = 2 × 10⁻⁴, also showing nominally significant group-by-sex interactions), increased relative slow power during N3 in frontal channels (F3: *b* = 0.77, *p* = 5 × 10⁻⁴), and greater SS chirp across all channels (C4: *b* = 0.56, *p* = 0.001).

### No sex differences in brain age prediction

Following the framework from our prior work (Kozhemiako et al., 2023), we also examined whether predicted age based on sleep EEG differed between boys and girls (**Figure 7A**). There were no significant sex differences in prediction accuracy, as measured by either mean absolute error or mean error, within any subgroup. However, both boys and girls with IntDis showed less precise predictions compared to their same-sex peers in the non-NDD group, as indicated by higher mean absolute error (girls: *b* = 0.96, *p* = 2 × 10⁻⁴; boys: *b* = 0.83, *p* = 0.003; **Figure 7B**). Additionally, boys were consistently predicted to be younger than their actual age based on mean error (*b* = –0.62, *p* = 0.01), with a similar trend observed in girls (*b* = –0.41, *p* = 0.09; **Figure 7C**).

**Figure 7.**
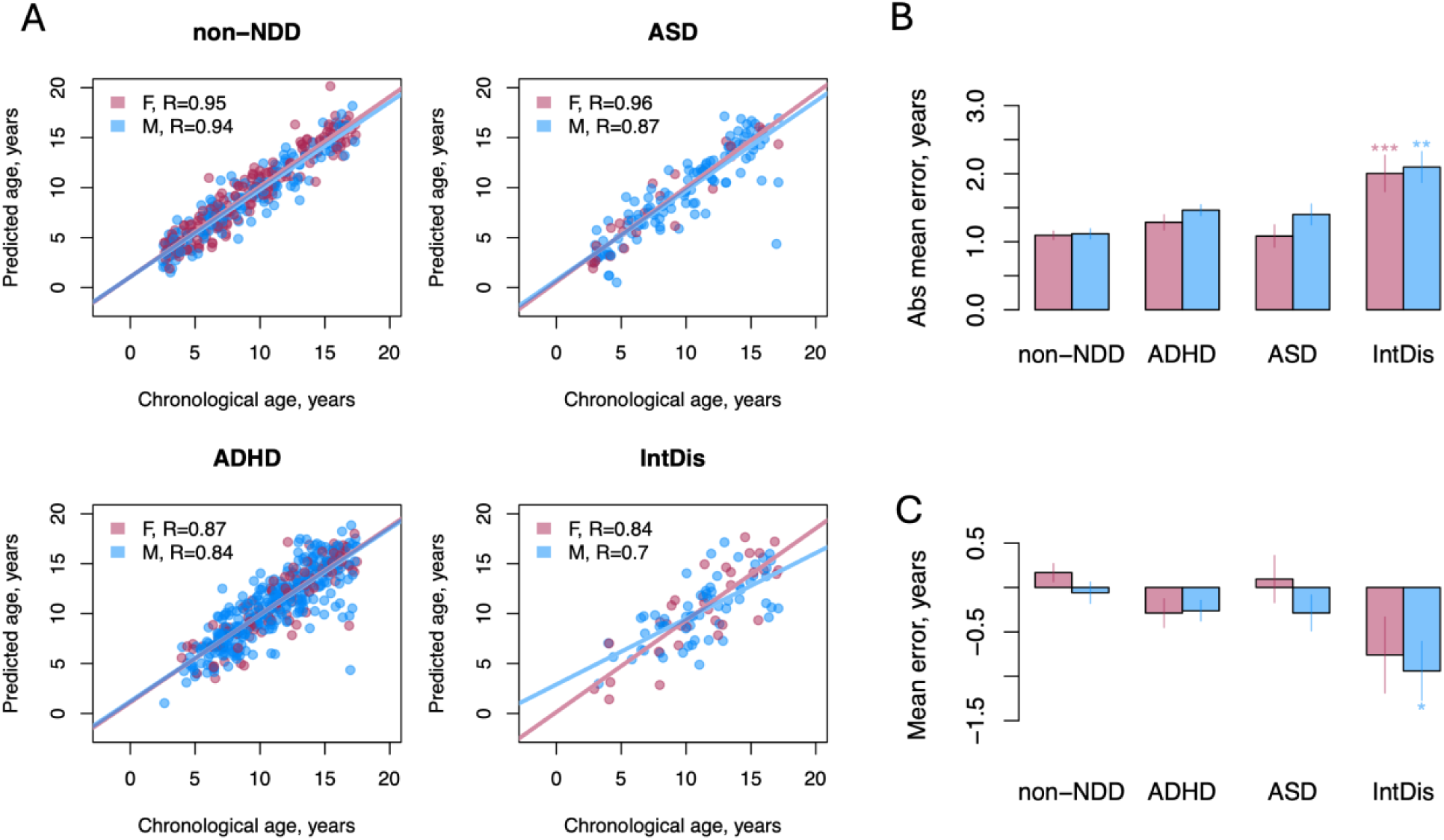
Brain age prediction in non-NDD and NDD children. A – the scatterplots illustrate correlation between the sleep EEG-predicted age and chronological age separately in boys and girls. B – the bar plot shows averaged absolute error in age prediction across sexes and groups with standard errors. C – the bar plot shows averaged error in age prediction across sexes and groups with standard errors. * – p < 0.05; ** – p < 0.01; *** – p < 0.001

## Discussion

In this study, we used a large clinical pediatric sample to investigate sex differences in both the prevalence of sleep disorders and sleep neurophysiology, as indexed by a broad range of EEG metrics. Sex differences in sleep EEG were also replicated in an independent cohort. To our knowledge, this is the largest study to date examining sex differences in sleep EEG features in non-NDD children, as well as in boys and girls with ADHD, ASD, and IntDis. Due to the strong male bias in NDD prevalence, it is typically challenging to collect sufficiently large samples to study girls with NDDs as a separate group. This makes large retrospective clinical datasets, such as the one from NCH, a valuable resource for research on sex differences in neurodevelopmental sleep physiology.

### Sex differences in sleep diagnoses

Since the cohort in this study consists of children referred for overnight sleep PSGs, the overall prevalence of sleep disorders is much higher than in the general population. In fact, over 80% of children in the non-NDD group had at least one sleep-related diagnosis documented in their medical records. This context is important when interpreting the findings. For example, previous studies have reported a strong association between obstructive sleep apnea (OSA) and ADHD, with up to 95% of children with OSA showing attention deficits (Youssef et al., 2011) and higher rates of ADHD diagnoses among children with OSA (Constantin et al., 2015). Yet, in our sample, both boys and girls with ADHD had significantly lower rates of OSA compared to non-NDD children – highlighting that the non-NDD group, drawn from a sleep clinic population, is likely oversaturated with sleep disorders relative to the general population.

Despite this elevated baseline, children in the NDD subgroups still showed significantly higher rates of certain sleep-related diagnoses, such as insomnia, compared to their non-NDD peers, regardless of sex. This aligns with prior research reporting increased insomnia in children with NDDs (Miano & Ferri, 2010; Wynchank et al., 2017) and further clarifies that these patterns are also present in girls when examined separately.

Within subgroups, sex differences were observed in the ADHD group, where boys had higher rates of OSA than girls – reflecting patterns seen in the general population (Redline et al., 2007). While no sex differences were found in the full non-NDD group (note that non-NDD group was on average younger than ADHD), a similar trend of increased rates of OSA in boys emerged when analyses were limited to older children in the NCH cohort (>12 years), supporting the idea that such differences may appear after puberty (Inoshita et al., 2018; Kim & Taranto-Montemurro, 2019).

No significant sex differences were observed within the IntDis and ASD group. Previous studies using parental reports in ASD summarized in the recent review have shown mixed results, with some indicating more sleep problems in boys, others in girls, and some showing no differences (Elkhatib Smidt et al., 2021).

### Differences in Sleep EEG

Our findings revealed multiple sex differences in sleep EEG metrics among non-NDD children. Importantly, despite the clinical nature of the NCH dataset, the sex effects were very similar to those in an independent, non-clinical sample. Boys generally exhibited more disrupted sleep macroarchitecture, characterized by higher sleep fragmentation, fewer sleep cycles, and shorter N2 duration. These results are consistent with prior PSG studies – albeit with smaller samples (fewer than 100 children) – which reported greater wake time after sleep onset (Markovic et al., 2020), shorter time in bed (Campbell et al., 2005) and lower percentage of slow wave sleep (Ringli et al., 2013) in school-age boys compared to girls, though some studies have found no significant sex differences (Baker et al., 2012).

There is strong evidence that good quality sleep is essential for healthy development (Kurth et al., 2015). Speculatively, the pattern of disrupted sleep observed in non-NDD boys may contribute to their increased vulnerability to neurodevelopmental alterations in childhood putting them at higher risk of NDDs. Interestingly, sex differences in sleep fragmentation measures were not evident in the NDD groups, where sleep macroarchitecture was generally similar between boys and girls with ADHD, ASD, or IntDis.

In contrast, some sleep microarchitecture metrics displayed consistent sex differences across all subsamples. For example, FS density was lower in boys compared to girls in the non-NDD, ADHD, ASD, and IntDis groups (although only nominally significant in the last two). This pattern aligns with findings from a previous study in a smaller non-NDD pediatric sample (Markovic et al., 2020) and appears to persist across the lifespan, as shown in a large mostly adult typical sample of over 10,000 individuals (Purcell et al., 2017). Additionally, in non-NDD group specifically, boys also exhibited lower amplitude and duration of FS, lower SS frequency, reduced sigma power and weaker FS coupling with SOs compared to girls. Many of these features are known to increase with age (Kozhemiako et al., 2023) and have been positively associated with cognitive performance in children (Reynolds et al., 2018).

Our findings suggest that higher values of these sleep metrics in girls may contribute to their relative resilience to neurodevelopmental disturbances during childhood.

Numerous sleep EEG metrics showed significant sex-by-group interactions across all NDDs. Notably, these effects were driven by greater alterations in girls than boys with ASD and ADHD when compared to same-sex non-NDD peers which may be consistent with the female protective effect theory. This theory suggests that girls who develop an NDD carry a higher underlying risk load; if disrupted sleep contributes to this load, our finding of more pronounced sleep alterations in girls aligns with this idea. Alternatively, if sleep disruptions are primarily a consequence of an NDD rather than part of the liability, greater disruptions in females could reflect diagnostic bias (Cruz et al., 2025). In other words, girls may need to exhibit more severe symptoms which were previously associated with more sleep issues (Tudor et al., 2012; Veatch et al., 2017) to receive a diagnosis. However, this was not the case in individuals with IntDis where boys exhibited larger alterations in sleep EEG compared to girls highlighting that the female protective effect hypothesis or diagnostic bias may not apply to all neurodevelopmental conditions exhibiting male preponderance.

We found sex-specific differences in spindle characteristics among children with ASD and IntDis compared to their same-sex non-NDD peers. In boys with ASD or IntDis, spindle amplitude was generally higher than in non-NDD boys, a pattern not significant in girls. Instead, girls with ASD or IntDis showed differences in spindle frequency. In the IntDis group specifically, spindle deficits varied by spindle type: boys had lower SS density, while girls had decreased FS density.

Previous studies of spindle alterations in NDDs have shown inconsistent results (Gruber & Wise, 2016). For example, reduced spindle density and duration were reported in preschool children with ASD (Farmer et al., 2018), but these findings were not replicated in the same extended sample using an automated spindle detection algorithm (Cumming et al., 2024). Instead, that study found increased intra-spindle frequency deceleration (or chirp) in ASD children compared to controls. Among older children, there are reports of higher spindle ratio between N3 vs N2 compared to controls (Kawahara et al., 2022), spindle reduction in ASD specific to N3 sleep (Mylonas et al., 2022), or no difference (Kurz et al., 2021). Notably, both of those latter studies included only boys. Research on spindles in children with IntDis is limited with just a few relatively older studies describing a range of abnormalities, including extremely high-amplitude spindles, absent spindles, or unusually short ones (Gruber & Wise, 2016). Our findings underscore that spindle alterations might differ between boys and girls, highlighting the importance of considering sex in future research on sleep neurophysiology in NDDs.

In contrast, R sleep alterations in ADHD and IntDis were consistent across sexes. Both boys and girls had longer R latency and, in case of IntDis group, shorter R stage duration. R sleep is believed to support crucial neurodevelopmental processes including synaptic pruning and memory consolidation (Li et al., 2017) and sensorimotor system development (Blumberg et al., 2013). It is also associated with dreaming, although its exact functions remain largely unclear (Tsunematsu, 2023). Notably, longer R duration has been linked to better cognitive outcomes in older adults (Djonlagic et al., 2020).

While data on R sleep in individuals with IntDis are limited, several studies report significantly reduced R sleep, consistent with our findings (Donnelly et al., 2022; Esposito & Carotenuto, 2014; Miano et al., 2008). In ADHD, the literature is more extensive but highly variable, ranging from reports of no differences in R latency (Kirov et al., 2004; Miano et al., 2019; Prehn-Kristensen et al., 2011; Virring et al., 2016), to shorter (Akinci et al., 2015; Darchia et al., 2021, 2022; Kirov et al., 2012) or longer latencies in ADHD (Castelnovo et al., 2022). Findings on R duration also included increased (Akinci et al., 2015; Kirov et al., 2004, 2012; Prehn-Kristensen et al., 2011; Virring et al., 2016), unchanged (Castelnovo et al., 2022; Darchia et al., 2021, 2022; Furrer et al., 2019; Miano et al., 2019; Prehn-Kristensen et al., 2013) or slightly reduced duration in ADHD (Saletin et al., 2017).

One possible reason for this inconsistency is the small sample sizes in previous studies – most included fewer than 30 ADHD participants, with only two exceptions [N = 76 (Virring et al., 2016), N = 50 (Furrer et al., 2019)]. In our large-scale analysis (N=525), we observed no difference in R duration but increased R latency in both boys and girls with ADHD. Importantly, a key distinction between our study and many previous ones is medication status. While we statistically controlled for psychotropic and CNS-active medications, most prior studies either used medication-naive participants or required a medication pause (typically 48 hours before sleep recording). Given that ADHD medications are known to affect sleep architecture (Kidwell et al., 2015) and R measures in particular (Sobanski et al., 2008), this may impact the comparability between our findings and earlier reports.

Although prior studies in older adults have shown significant sex differences in brain age estimated from sleep EEG with males exhibiting older predicted brain age than females (Brink-Kjaer et al., 2022), we found no such difference in our pediatric sample. Brain age predictions were comparable between boys and girls, regardless of diagnostic status. This suggests that despite some sex-specific patterns, age-related changes in sleep EEG follow broadly similar developmental trajectories in boys and girls during childhood. We also found that previously reported less precise age prediction for children with IntDis compared to non-NDD children (Kozhemiako et al., 2023) was observed in both boys and girls with IntDis.

Overall, our findings indicate that sleep EEG exhibits multiple sex-specific patterns during childhood in the non-NDD population, many of which are altered in the NDD children. We observed more disrupted sleep and fewer FS in boys. Given the role of FS in cognition and brain maturation, this finding may partly explain boys’ greater vulnerability to neurodevelopmental perturbations in childhood compared to girls. Despite these sex differences in individual sleep metrics, the composite brain age prediction estimates did not differ between boys and girls suggesting that overall childhood developmental trajectories of sleep EEG are broadly similar across sexes. However, many EEG alterations in NDDs relative to non-NDD children varied between boys and girls, emphasizing the importance of including sex as a biological variable in future research. These differences may reflect varying levels of sleep disruption or be linked to disorder-relevant factors such as differences in symptom expression. Future studies should align symptom profiles with sleep EEG measures to clarify these relationships.

