## Supplementary Table S1 for "The effect of sex on sleep EEG in typical and altered neurodevelopment"

### SUPPLEMENTARY MATERIAL

#### Supplementary table 1

| Disorders | N | Diagnosis name | Diagnosis ICD codes |
|---|---|---|---|
| ASD | 196 | Autistic disorder; Autistic disorder, current or active state | F84.0; 299.00 |
| ADHD | 525 | Attention-deficit hyperactivity disorder, combined type; Attention deficit disorder with hyperactivity(314.01); Attention-deficit hyperactivity disorder, unspecified type; Attention-deficit hyperactivity disorder, predominantly inattentive type; Attention-deficit hyperactivity disorder, other type; Attention deficit disorder without mention of hyperactivity; Attention-deficit hyperactivity disorder, predominantly hyperactive type | F90.2; 314.01; F90.9; F90.0; F90.8; 314.00; F90.1 |
| IntDis | 167 | Unspecified intellectual disabilities; Moderate intellectual disabilities; Mild intellectual disabilities; Severe intellectual disabilities; Profound intellectual disabilities; Other intellectual disabilities | F79; F71; F70; F72; 319; 318.2; 317; F78; 318.1; F73; 318.0 |
| DS | 140 | Down syndrome, unspecified; Down's syndrome; Trisomy 21, mosaicism (mitotic nondisjunction); Trisomy 21, translocation; Trisomy 21, nonmosaicism (meiotic nondisjunction) | Q90.9; 758.0; Q90.1; Q90.2; Q90.0 |
| CP | 138 | Spastic quadriplegic cerebral palsy; Cerebral palsy, unspecified; Other cerebral palsy; Spastic diplegic cerebral palsy; Spastic hemiplegic cerebral palsy; Infantile cerebral palsy, unspecified; Other specified infantile cerebral palsy; Athetoid cerebral palsy; Ataxic cerebral palsy | G80.0; G80.9; G80.8; G80.1; G80.2; 343.9; 343.8; G80.3; 333.71; G80.4 |
| Epilepsy | 242 | Epilepsy, unspecified, not intractable, without status epilepticus; Localization-related (focal) (partial) symptomatic epilepsy and epileptic syndromes with complex partial seizures, not intractable, with status epilepticus; Generalized idiopathic epilepsy and epileptic syndromes, not intractable, without status epilepticus; Localization-related (focal) (partial) symptomatic epilepsy and epileptic syndromes with complex partial seizures, not intractable, without status epilepticus; Epileptic spasms, not intractable, without status epilepticus; Localization-related (focal) (partial) symptomatic epilepsy and epileptic syndromes with simple partial seizures, not intractable, without status epilepticus; Todd's paralysis (postepileptic); Lennox-Gastaut syndrome, intractable, with status epilepticus; Localization-related (focal) (partial) symptomatic epilepsy and epileptic syndromes with complex partial seizures, intractable, without status epilepticus; Generalized idiopathic epilepsy and epileptic syndromes, intractable, without status epilepticus; Lennox-Gastaut syndrome, intractable, without status epilepticus; Absence epileptic syndrome, not intractable, without status epilepticus; Other generalized epilepsy and epileptic syndromes, not intractable, without status epilepticus; Absence epileptic syndrome, intractable, without status epilepticus; Other generalized epilepsy and epileptic syndromes, intractable, without status epilepticus; Localization-related (focal) (partial) symptomatic epilepsy and epileptic syndromes with simple partial seizures, intractable, without status epilepticus; Epilepsy, unspecified, intractable, without status epilepticus; Other epilepsy, intractable, without status epilepticus; Lennox-Gastaut syndrome, not intractable, without status epilepticus; Epileptic spasms, intractable, with status epilepticus; Localization-related (focal) (partial) symptomatic epilepsy and epileptic syndromes with complex partial seizures, intractable, with status epilepticus; Localization-related (focal) (partial) idiopathic epilepsy and epileptic syndromes with seizures of localized onset, intractable, without status epilepticus; Epilepsy, unspecified, not intractable, with status epilepticus; Other epilepsy, not intractable, without status epilepticus; Epileptic spasms, intractable, without status epilepticus; Unspecified epilepsy without mention of intractable epilepsy; Epilepsy, unspecified, intractable, with status epilepticus; Localization-related (focal) (partial) epilepsy and epileptic syndromes with complex partial seizures, without mention of intractable epilepsy; Infantile spasms with intractable epilepsy; Other epilepsy, not intractable, with status epilepticus; Localization-related (focal) (partial) idiopathic epilepsy and epileptic syndromes with seizures of localized onset, not intractable, without status epilepticus; Localization-related (focal) (partial) idiopathic epilepsy and epileptic syndromes with seizures of localized onset, not intractable, with status epilepticus; Infantile spasms without mention of intractable epilepsy; Unspecified epilepsy with intractable epilepsy; Localization-related (focal) (partial) epilepsy and epileptic syndromes with simple partial seizures, without mention of intractable epilepsy; Generalized nonconvulsive epilepsy without mention of intractable epilepsy; Other forms of epilepsy and recurrent seizures without mention of intractable epilepsy; Generalized convulsive epilepsy without mention of intractable epilepsy; Generalized convulsive epilepsy with intractable epilepsy; Localization-related (focal) (partial) epilepsy and epileptic syndromes with complex partial seizures, with intractable epilepsy; Generalized nonconvulsive epilepsy with intractable epilepsy; Epileptic grand mal status; Generalized idiopathic epilepsy and epileptic syndromes, not intractable, with status epilepticus; Other generalized epilepsy and epileptic syndromes, not intractable, with status epilepticus; Other generalized epilepsy and epileptic syndromes, intractable, with status epilepticus; Epileptic petit mal status; Localization-related (focal) (partial) symptomatic epilepsy and epileptic syndromes with simple partial seizures, not intractable, with status epilepticus; Epileptic seizures related to external causes, not intractable, with status epilepticus; Other epilepsy, intractable, with status epilepticus; Localization-related (focal) (partial) symptomatic epilepsy and epileptic syndromes with simple partial seizures, intractable, with status epilepticus; Generalized idiopathic epilepsy and epileptic syndromes, intractable, with status epilepticus; Lennox-Gastaut syndrome, not intractable, with status epilepticus; Localization-related (focal) (partial) epilepsy and epileptic syndromes with simple partial seizures, with intractable epilepsy; Juvenile myoclonic epilepsy, intractable, without status epilepticus; Other forms of epilepsy and recurrent seizures with intractable epilepsy; Absence epileptic syndrome, not intractable, with status epilepticus; Epileptic seizures related to external causes, not intractable, without status epilepticus; Localization-related (focal) (partial) idiopathic epilepsy and epileptic syndromes with seizures of localized onset, intractable, with status epilepticus; Epileptic spasms, not intractable, with status epilepticus | G40.909; G40.201; G40.309; G40.209; G40.822; G40.109; G83.84; G40.813; G40.219; G40.319; G40.814; G40.A09; G40.409; G40.A19; G40.419; G40.119; G40.919; G40.804; G40.812; G40.823; G40.211; G40.019; G40.901; G40.802; G40.824; 345.90; G40.911; 345.40; 345.61; G40.801; G40.009; G40.001; 345.60; 345.91; 345.50; 345.00; 345.80; 345.10; 345.11; 345.41; 345.01; 345.3; G40.301; G40.401; G40.411; 345.2; G40.101; G40.501; G40.803; G40.111; G40.311; G40.811; 345.51; G40.B19; 345.81; G40.A01; G40.509; G40.011; G40.821 |
| Language disorders | 783 | Mixed receptive-expressive language disorder; Other developmental disorders of speech and language; Developmental disorder of speech and language, unspecified; Expressive language disorder; Other developmental speech or language disorder; Care involving speech-language therapy | F80.2; F80.89; F80.9; F80.1; 315.39; V57.3; 315.32; 315.31 |
| Tic disorders | 19 | Other tic disorders; Chronic motor or vocal tic disorder; Transient tic disorder | F95.8; F95.1; 307.21; F95.0; 307.22 |
| Central apnea | 58 | Primary central sleep apnea; Central sleep apnea in conditions classified elsewhere(327.27); Central sleep apnea in conditions classified elsewhere | G47.31; 327.21; 327.27; G47.37 |
| Other apneas | 2008 | Obstructive sleep apnea (adult) (pediatric); Sleep apnea, unspecified; Apnea, not elsewhere classified; Other sleep apnea; Unspecified sleep apnea; Apnea; Primary apnea of newborn; Other apnea of newborn; Other organic sleep apnea; Organic sleep apnea, unspecified | G47.33; G47.30; R06.81; G47.39; 327.23; 780.57; 786.03; 770.81; P28.4; 770.82; 327.29; 327.20 |
| Insomnia | 403 | Insomnia due to medical condition; Insomnia due to other mental disorder; Insomnia, unspecified; Other insomnia; Problems related to behavioral insomnia of childhood; Psychophysiologic insomnia; Behavioral insomnia of childhood, sleep-onset association type; Behavioral insomnia of childhood, unspecified type; Behavioral insomnia of childhood, combined type; Primary insomnia; Behavioral insomnia of childhood, limit setting type; Insomnia due to medical condition classified elsewhere; Other insomnia not due to a substance or known physiological condition; Adjustment insomnia | G47.01; F51.05; G47.00; G47.09; 780.52; V69.5; F51.04; Z73.810; Z73.819; Z73.812; F51.01; Z73.811; 327.01; F51.09; F51.02 |

| | | | |
|---|---|---|---|
| Hypersomnia | 175 | Hypersomnia, unspecified; Other hypersomnia; Idiopathic hypersomnia with long sleep time; Hypersomnia due to medical condition; Idiopathic hypersomnia without long sleep time | G47.10; G47.19; G47.11; 780.54; G47.14; G47.12 |
| Circadian rhythm disorders | 440 | Circadian rhythm sleep disorder, unspecified type; Circadian rhythm sleep disorder, irregular sleep wake type; Circadian rhythm sleep disorder, unspecified Circadian rhythm sleep disorder of nonorganic origin; Circadian rhythm sleep disorder, free running type; Circadian rhythm sleep disorder, advanced sleep phase type | G47.20, G47.23, 327.30; G47.21, 307.45; G47.24; G47.22 |
| Other sleep disorders | 845 | Sleep disorder, unspecified; Sleep terrors (night terrors);<br>Restless legs syndrome; Sleep deprivation; Other sleep disorders; Other sleep disturbances; Sleep disturbance, unspecified; Restless legs syndrome (RLS); Narcolepsy with cataplexy; Idiopathic sleep related nonobstructive alveolar hypoventilation; Parasomnia, unspecified; Sleep related hypoventilation/hypoxemia in conditions classifiable elsewhere; Problems related to lack of adequate sleep; Sleep related bruxism; Sleepwalking (somnambulism); Dysfunctions associated with sleep stages or arousal from sleep, Other dysfunctions of sleep stages or arousal from sleep; Other specific disorder of sleep of nonorganic origin; Sleep arousal disorder; Inadequate sleep hygiene; Other psychoactive substance use, unspecified with psychoactive substance-induced sleep disorder; Other sleep related movement disorders; Other sleep disorders not due to a substance or known physiological condition; Other parasomnia 16<br>Persistent disorder of initiating or maintaining sleep; Sleep related hypoventilation in conditions classified elsewhere; Narcolepsy without cataplexy; Sleep disorder not due to a substance or known physiological condition, unspecified; Insufficient sleep syndrome; Narcolepsy in conditions classified elsewhere without cataplexy; Sleep related movement disorder, unspecified; Recurrent isolated sleep paralysis; REM sleep behavior disorder; Parasomnia in conditions classified elsewhere; Other organic sleep disorders; Nonorganic sleep disorder, unspecified; Narcolepsy without cataplexy(347.00); Sleep related leg cramps | G47.9; F51.4; G25.81; Z72.820; G47.8; 780.59; 780.50; 333.94; G47.411; G47.34; G47.50; 327.26; V69.4; G47.63; F51.3; 780.56; 307.47; 307.49; 307.46; Z72.821; F19.982; G47.69; F51.8; G47.59; 307.42; G47.36; G47.419; F51.9; F51.12; 327.53; G47.429; 780.58; G47.53; G47.52; G47.54; 327.8; 307.40; 327.24; 327.42; 347.00; G47.62 |

Supplementary table 2 Medication overlapping PSG recording in NCH sample

| Therapeutical class | Pharmaceutic class | Therapeutic subclass |
|---|---|---|
| ANTIHISTAMINES | | |
| | ANTIHISTAMINES - 2ND GENERATION | cetirizine HCl; loratadine; fexofenadine HCl |
| | ANTIHISTAMINES - 1ST GENERATION | hydroxyzine HCl; cyproheptadine HCl; diphenhydramine HCl; ; hydro xyzine pamoate; promethazine HCl |
| | EYE ANTIHISTAMINES | azelastine HCl; ketotifen fumarate |
| PSYCHOTHERAPEUTIC DRUGS | | |
| | SELECTIVE SEROTONIN REUPTAKE INHIBITOR (SSRIS) | citalopram hydrobromide; fluoxetine HCl; escitalopram oxalate; sert raline HCl; fluvoxamine maleate |
| | ALPHA-2 RECEPTOR ANTAGONIST ANTIDEPRESSANTS | mirtazapine |
| | ANTIPSYCHOTIC,ATYPICAL,DOPAMINE,SEROTONIN ANTAGNST | risperidone; quetiapine fumarate; lurasidone HCl; olanzapine |
| | TRICYCLIC ANTIDEPRESSANTS,REL.NON-SEL.REUPT-INHIB | amitriptyline HCl; doxepin HCl; ; clomipramine HCl |
| | ANTIPSYCHOTICS, ATYP, D2 PARTIAL AGONIST/5HT MIXED | aripiprazole |
| | TX FOR ADHD - SELECTIVE ALPHA-2 RECEPTOR AGONIST | guanfacine HCl; clonidine HCl |
| | SEROTONIN-2 ANTAGONIST/REUPTAKE INHIBITORS (SARIS) | trazodone HCl |
| | NARCOLEPSY AND SLEEP DISORDER THERAPY AGENTS | modafinil; armodafinil |
| | TX FOR ATTENTION DEFICIT-HYPERACT(ADHD)/NARCOLEPSY | methylphenidate HCl; dexmethylphenidate HCl |
| | ANTI-ANXIETY - BENZODIAZEPINES | diazepam; lorazepam |
| | TX FOR ATTENTION DEFICIT-HYPERACT.(ADHD), NRI-TYPE | atomoxetine HCl |
| | ADRENERGICS, AROMATIC, NON-CATECHOLAMINE | lisdexamfetamine dimesylate |
| | ANTI-ANXIETY DRUGS | buspirone HCl |
| | BIPOLAR DISORDER DRUGS | lithium carbonate |
| | SEROTONIN-NOREPINEPHRINE REUPTAKE-INHIB (SNRIS) | venlafaxine HCl; duloxetine HCl |
| | NOREPINEPHRINE AND DOPAMINE REUPTAKE INHIB (NDRIS) | bupropion HCl |
| CNS DRUGS | | |
| | ANTICONVULSANT - BENZODIAZEPINE TYPE | clobazam; diazepam; clonazepam |
| | ANTICONVULSANTS | divalproex sodium; zonisamide; oxcarbazepine; valproic acid (as sodi um salt); ; topiramate; rufinamide; lacosamide; pregabalin; gabapen tin; lamotrigine; brivaracetam; ethosuximide; levetiracetam; felbam ate; vigabatrin; carbamazepine; phenytoin; perampanel; diazepam |
| | ANTICONVULSANT - CANNABINOID TYPE | cannabidiol (CBD) |
| HORMONES | | |
| | GROWTH HORMONES | somatropin |
| | PINEAL HORMONE AGENTS | melatonin; |

| | | |
|---|---|---|
| | GLUCOCORTICOIDS | prednisone; hydrocortisone; hydrocortisone sodium succ/PF; predni solone sodium phosphate; hydrocortisone sod succinate; deflazacort; methylprednisolone; dexamethasone sodium phosphat e |
| | ANTIDIURETIC AND VASOPRESSOR HORMONES | desmopressin acetate |
| | PROGESTATIONAL AGENTS | norethindrone acetate; medroxyprogesterone acetate |
| | LHRH(GNRH)AGNST PIT.SUP-CENTRAL PRECOCIOUS PUBERTY | leuprolide acetate |
| | MINERALOCORTICOIDS | fludrocortisone acetate |
| | ESTROGENIC AGENTS | estradiol |
| SEDATIVE/HYPNOTICS | | |
| | BARBITURATES | phenobarbital |
| | SEDATIVE-HYPNOTICS - BENZODIAZEPINES | lorazepam |

**Table S3 Demographic characteristics of the samples in replication analysis**

| **Characteristics** | **NCH subsample** | | **CHAT** | |
|---|---|---|---|---|
| | **Non-NDD boys** | **Non-NDD girls** | **Boys** | **Girls** |
| **N** | 662 | 499 | 571 | 630 |
| **Age, M (SD) age range, years** | 7.2 (1.6) 4.5 - 10 | 7 (1.5) 4.5 - 10 | 7 (1.4) 4.5 - 10 | 7.1 (1.4) 4.8 - 10 |
| **Self-reported race: Whites** | 466 (70%) | 335 (67%) | 243 (44%) | 239 (38%) |
| **Blacks** | 108 (16%) | 89 (18%) | 253 (43%) | 312 (50%) |
| **Other** | 66 (10%) | 62 (12%) | 70 (12%) | 70 (12%) |
| **Unknown** | 22 (3%) | 13 (3%) | 0 (0%) | 0 (0%) |
